# GRASER: haplotype assignment of graph-aligned short reads

**DOI:** 10.64898/2026.11.13.766454

**Authors:** John E. Garza, Juan F. Macias-Velasco, Derek A. Albracht, Edward A. Belter, Chad Tomlinson, Daofeng Li, Wenjin Zhang, Ting Wang

## Abstract

Existing read phasing tools operate on reads aligned to a linear reference, where reference bias reduces the information available for haplotype inference and read assignment. Pangenome graphs reduce reference bias, but reads falling within non-reference sequence cannot be surjected to a linear reference, so phasing after surjection reintroduces that bias. Here we present GRASER, which phases graph-aligned short reads against haplotype paths inferred for each individual, using established graph genotyping and statistical phasing to build those paths. GRASER phase uses edges unique to one haplotype path as evidence, so reads are phased directly from graph alignments without conversion to a linear reference. Against true parental haplotype paths, GRASER assigned 22.6 to 24.5% of simulated fragments to a haplotype, and more than 96% of those were placed on the correct haplotype. Inferred haplotype paths reduced the assignment rate by about 4 percentage points. GRASER assigned more fragments than a WhatsHap-based linear-reference workflow in every configuration tested. Applied to whole-genome bisulfite sequencing, GRASER recovered allele-specific methylation at imprinted loci and measured haplotype-resolved methylation within a structural variant.

## INTRODUCTION

Humans are diploid, carrying two copies of each autosome, one inherited from each parent. Where the two copies differ in sequence, each alternative form at a locus is an allele, and the collection of alleles carried on a single chromosome copy is a haplotype. A cis-genetic effect is a difference in molecular phenotype, such as gene expression or DNA methylation, that a variant produces only on the haplotype that carries it^1,3^. Because both haplotypes share the same cellular environment, comparing them within an individual separates cis effects from trans-acting influences. Identifying these effects therefore requires phasing, of both variants and sequencing reads.

Variant phasing determines which alleles across heterozygous sites lie together on the same haplotype. Statistical phasing, as implemented in SHAPEIT4^5^, models each haplotype in a sample as a mosaic of haplotypes observed in a population, using linkage disequilibrium and recombination to infer the phase of observed genotypes. These population haplotypes may come from the study cohort or from an external reference panel. Because phase is inferred from the population rather than observed in the reads, statistical phasing can establish long-range phase relationships without requiring reads to span variants, making it well suited to short-read sequencing and genotyping array data. However, its accuracy depends on the availability of informative population haplotypes and may be reduced for rare variants or for populations poorly represented in the reference panel.

Another approach is read-informed phasing, implemented by tools such as WhatsHap phase^2^. Reads spanning at least two heterozygous variants are used to link the observed alleles to the same haplotype. Overlapping reads can be used to chain co-occurring variants together into larger regions known as phase blocks. Since phase evidence is derived from the sample being analyzed, allele assignment accuracy is not subject to population frequency bias. Instead, this technique is limited by read depth, read length, and the density of heterozygous variants. Short read data often cannot provide sufficient information to link distant variants, resulting in fragmented phase blocks.

Once variants have been phased, they can be used to assign reads to their haplotype of origin. This is known as read phasing or haplotagging, and is implemented by tools such as WhatsHap haplotag^2^. Alleles observed in aligned reads are compared to known phased variants to determine which haplotype they support. Separating reads by haplotype enables allele-specific quantification of molecular phenotypes, such as gene expression, DNA methylation, and chromatin accessibility, providing evidence for cis-genetic effects. However, read phasing is also limited by read length. Because distinguishing heterozygous sites are sparse across the genome, only a small proportion of short reads can be confidently assigned to a haplotype.

Read phasing is also affected by the reference used for alignment. Conventional analyses use a linear reference, which contains a single haploid sequence. A linear reference cannot represent the alternative haplotypes present within an individual or across a population. Reads carrying non-reference alleles align less accurately, receive lower mapping quality, or fail to align^4,6^. This overrepresents reference alleles among aligned reads, an effect known as reference bias. Reference bias reduces the number of variants identified, and with it the information available for haplotype inference and read assignment. Because the reads that remain favor the reference allele, reference bias also skews observed allele-specific measurements^1,3,4,7^. Personalized references that incorporate known variants reduce this bias. However, those variants are called from alignments to the linear reference, so the personalized reference retains part of the bias it is meant to correct.

Pangenome graphs offer an alternative to linear references by encoding the variation present across many individuals^8,9^. Sequence is stored in nodes, and edges connect adjacent sequences. Alternative alleles are represented as different traversals through a locus, with individual haplotypes as paths through the graph. Because the graph incorporates variation from many haplotypes, reads from both haplotypes of an individual align with similar accuracy, which reduces reference bias^6,10^.

Existing read phasing tools operate on reads aligned to a linear reference. Graph alignments can be converted to linear alignments by surjection, which maps alignments from all paths in the graph onto a single reference path^6,8,11^. Reads falling within non-reference sequence, such as an insertion, have no syntenic position on the reference path and cannot be surjected. Because these reads come from the non-reference allele, their loss reintroduces reference bias. Taking full advantage of graph alignment therefore requires read phasing performed directly on the graph.

Here we present GRASER, a graph-native tool that assigns graph-aligned short reads to haplotypes. The GRASER path module uses genome inference to build personalized haplotype paths through the graph, combining graph-based genotyping of an individual’s reads with statistical phasing^5,12,13^. GRASER phase then assigns reads to one of two haplotype paths, either those built by GRASER path or paths already present in the graph. Because variation introduces node boundaries in the graph, edges unique to one haplotype span sequence that distinguishes the two. GRASER phase uses these haplotype-unique edges as evidence for assignment. We evaluate GRASER using whole-genome sequencing (WGS) from seven individuals from Genome in a Bottle and the 1000 Genomes Project, and compare it with a linear-reference workflow. To demonstrate its use with functional data, we apply GRASER to paired WGS and whole-genome bisulfite sequencing (WGBS) from EN-TEx to detect allele-specific methylation.

## MATERIALS & METHODS

### Sequencing Data & Resources

#### Sequencing Datasets & Preprocessing

We evaluated GRASER using WGS from seven individuals in two cohorts. To test whether performance depends on an individual’s assemblies being present in the reference graph, and to draw on extensive high-quality annotations, we selected HG002 and HG005 from Genome in a Bottle, forming the GIAB cohort. To assess performance across a broader set of samples, we added five individuals from the 1000 Genomes Project, forming the 1kGP cohort. To assess performance on real data, we downloaded WGS from NCBI SRA for HG002 (SRR14724532) and HG005 (SRR14724509). Measuring assignment accuracy and the effect of genomic context requires knowing each read’s haplotype and position of origin. We therefore simulated error-free 150 bp paired-end reads from both parental haplotype assemblies of each GIAB individual^9^ to 15X per haplotype. To sample both haplotypes evenly, so that phasing could be evaluated uniformly across the genome, we used the deterministic lattice simulator in MoRGANA. Reads from the two haplotypes were combined to produce 30X diploid data. The 1kGP cohort comprised HG00621 (ERX3990668), HG00741 (ERX3990691), HG01952 (ERX3990823), HG01978 (ERX3990830), and HG03516 (ERX3991039).

To demonstrate GRASER on functional data, we used paired WGS and WGBS from a single individual in the EN-TEx resource (donor ENCDO845WKR). Bisulfite treatment converts unmethylated cytosines to thymines, so C-to-T variants cannot be distinguished from conversion in WGBS reads. We therefore used the WGS to build a personalized graph containing phased haplotype paths inferred by GRASER path. The WGBS reads were then aligned to this graph with methylGrapher^10^. GRASER phase assigned the aligned reads to haplotypes, giving DNA methylation measurements for each donor haplotype. We used WGS library ENCLB480CRB and WGBS experiments from six tissues: spleen (ENCSR757JTX), lower leg skin (ENCSR128RMY), testis (ENCSR806NNG), thyroid (ENCSR403DTW), pancreas (ENCSR996BJT), and left lung upper lobe (ENCSR224CTR).

The EN-TEx WGS library was sequenced across three lanes, and the corresponding FASTQ files for each mate were concatenated before preprocessing. All other real libraries had a single FASTQ file per mate. All real data were preprocessed with fastp^14^ using the options ––dedup and ––detect_adapter_for_pe. Following the protocol used previously^10^, WGBS data were processed with the additional options ––trim_front1 10, ––trim_front2 15, ––cut_tail, ––cut_tail_mean_quality 20, ––qualified_quality_phred 20, and ––length_required 20.

#### Pangenome Reference Graphs

To assess whether GRASER’s performance depends on the evaluated individual’s assemblies being included in the reference pangenome, we compared results with and without them. We performed these paired analyses using two CHM13-based Minigraph-Cactus graphs from the second release of the Human Pangenome Reference Consortium (HPRC)^9^. The standard HPRCv2 graph comprises 231 pairs of haplotype assemblies, including all seven study individuals, in addition to GRCh38 and CHM13. In contrast, the HPRCv2 evaluation graph (HPRCv2-eval) was constructed with three individuals excluded, including both GIAB cohort individuals. Both graphs were used as references for read alignment and as sources of variation for personalized haplotype inference. GIAB cohort individuals were evaluated with both graphs.

Both graphs were built with the Minigraph-Cactus pangenome pipeline as part of the HPRC release^9,15^. We used the CHM13-based graphs because CHM13 is a complete, telomere-to-telomere assembly^16^, and a complete reference reduces reference bias^4^. The primary reference serves as the backbone of the graph and is the only assembly guaranteed to be preserved in its entirety. Secondary references, here GRCh38, are aligned next, followed by all other input assemblies in order of decreasing sequence similarity. Large unaligned sequences (>10 kb) are then clipped from all assemblies except the primary reference.

### GRASER Workflow

GRASER phase assigns graph-aligned reads to one of two haplotype paths. These paths may already be present in the graph, or may be inferred from an individual’s reads by GRASER path.

#### WGS Alignment

Aligning directly to the complete HPRC graph reduces performance, because variation absent from the target individual confounds alignment at many loci^12^. Current best practice is therefore to align to a personalized subgraph^12^. We counted k-mers in each preprocessed WGS library with KMC^17^. Given these counts, vg Giraffe^18^ runs in haplotype sampling mode^12^, building a diploid personalized graph (DPG) from the complete graph and then aligning reads to it. All WGS libraries were aligned to the HPRCv2 graph with default parameters. GIAB cohort libraries were also aligned to the HPRCv2-eval graph.

#### SHAPEIT4 Reference Data

Statistical phasing infers haplotypes from those observed across many individuals, and our seven individuals were too few to phase as a cohort, so we phased each against a population reference panel with SHAPEIT4^5^. We used a CHM13-based haplotype reference panel of 3,202 phased individuals from the 1000 Genomes Project^19^. This panel includes the 1kGP cohort individuals but not the GIAB cohort individuals. SHAPEIT4 also uses a recombination map to estimate where switches between panel haplotypes are likely, and we used the CHM13 recombination map from the same study^19^.

#### Variant Genotyping & Statistical Variant Phasing

For each alignment, we computed read support with vg pack and genotyped graph variants with vg call^13^, using the complete graph from which the personalized graph was sampled. Using bcftools^20^, each resulting VCF was normalized, filtered to remove non-PASS variants, and split by chromosome. Each VCF was phased with SHAPEIT4 using the reference panel and recombination map described above, producing a phased subset of the input variants. To preserve variants not phased by SHAPEIT4 while incorporating inferred phase from eligible sites, we used bcftools annotate to apply phased genotypes to the corresponding records in the input VCF.

#### Inferred Haplotype Path Construction (GRASER path)

GRASER path converts phased genotypes into a pair of haplotype paths through the graph. The phased VCF is exported to a tab-separated file with bcftools query. GRASER path takes this file and the name of a reference path as input. Because every site needs a phase for the paths to be complete, GRASER path assigns a random phase to each site left unphased by SHAPEIT4. We set a fixed seed so that this assignment is deterministic. For each haplotype, GRASER path then extracts the allele traversal at each site, which vg call records in the INFO/AT field as the path through the graph representing that allele. Because the first and last nodes of every traversal lie on the reference path, the gaps between variant sites are filled with reference sequence. We used CHM13 as the reference path, so the inferred haplotype paths are contiguous across each chromosome. GRASER path writes these paths as GFA W-lines.

#### Read Phasing (GRASER phase)

GRASER phase takes two inputs, haplotype paths and graph alignments. Paths are read from GFA W-lines, so either a pangenome graph or the output of GRASER path can be supplied. The target pair of paths is identified by sample name and haplotype indices. Alignments are supplied in GAF format. Both single-end and paired-end reads are supported, and paired-end alignments must be interleaved.

Sequence variation introduces node boundaries in the graph, so edges unique to one haplotype span sequence that distinguishes the pair. These haplotype-unique edges provide the evidence for read assignment. GRASER phase therefore first builds an index of the edges in the two target paths. Each path is broken into ordered pairs of oriented nodes. The edges of each haplotype are pooled across the whole genome in global scoring mode, or per contig in contig scoring mode. Each edge is then classified as shared, unique to haplotype 1, or unique to haplotype 2. A read that aligns within a single node traverses no edges, so GRASER phase also indexes haplotype-unique nodes and uses them to rescue these reads. Contig scoring mode requires reads to be assigned to a contig; since this is not always possible, all analyses used global scoring mode.

GRASER phase then processes the alignments one read or read pair at a time. Each edge in the read’s alignment traversal is weighted for each haplotype, in both the forward and reverse-complement orientations. Edges not present in the edge index are given a small non-zero weight ε (default 0.001), so that the logarithm in the score is always defined. Edges found in a haplotype are given a weight of 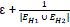, where the denominator is the number of distinct edges in the union of the two haplotype paths. Each haplotype’s score is the sum of the base-10 logarithms of its edge weights.

Because a read’s traversal may run in either orientation relative to the haplotype paths, the orientation that maximizes the score is selected for each haplotype. For paired-end reads, the mates are scored jointly in the two orientations consistent with a single fragment, mate 1 forward with mate 2 reverse-complemented or the reverse, and the combination with the higher combined score is selected. GRASER phase then computes a score ratio (SR) for each read by subtracting the haplotype 2 score from the haplotype 1 score. Because the scores are logarithms, this difference is the log ratio of the two haplotype scores. A read is assigned to haplotype 1 if SR > τ (default 1 × 10⁻⁶), to haplotype 2 if SR < −τ, and is otherwise called ambiguous.

Because both mates come from the same DNA fragment, their assignments should agree, so disagreement between mates indicates an unreliable assignment. If both mates are assigned to the same haplotype, the pair is labeled concordant and assigned to that haplotype. If they are assigned to opposite haplotypes, the pair is labeled discordant and left unassigned. If one mate is assigned and the other is ambiguous, the pair is labeled partially concordant and assigned to the informative mate’s haplotype. If both mates are ambiguous, the pair is labeled ambiguous. Finally, an ambiguous read or pair in which at least one mate aligns entirely within a single node is assigned to a haplotype if that node is in the haplotype-unique node index. If both mates are single node alignments and are assigned to different haplotypes, the fragment is left ambiguous.

GRASER phase can randomly assign ambiguous reads or pairs to a haplotype (––randomize-ambiguous), using a fixed seed for reproducibility. We enabled this option for WGS, but excluded randomly assigned reads from all evaluations. We did not enable it for WGBS, because randomly assigned reads would mix methylation signal from the two haplotypes. Without random assignment, only reads carrying evidence for a haplotype are assigned, restricting haplotype-resolved methylation to CpGs within read length of an informative variant.

### Performance Evaluation

#### Reference Datasets and Annotations

Evaluating genotyping and variant phasing requires each individual’s true phased genotypes, which we took from their haplotype assemblies in the HPRCv2 graph. We retrieved the CHM13-based VCF released with the standard HPRCv2 graph, which contains phased genotypes for every assembly in the graph, extracted with vg deconstruct by the Minigraph-Cactus pipeline. Using bcftools, the VCF was normalized, split by chromosome, and subset to each evaluated individual. These per-individual VCFs served as truth sets.

To assess how genomic context affects read phasing, we annotated each simulated fragment by intersecting its known location of origin with two features. To capture genomic complexity, we used regions annotated as difficult on both HG002 parental haplotype assemblies by Genome in a Bottle. Fragments in which at least one mate overlapped these regions were labeled difficult. To identify fragments likely to carry haplotype-distinguishing variation, we called high-confidence heterozygous sites between the two parental assemblies of HG002 and HG005 with Dipcall^21^. Fragments in which at least one mate overlapped a Dipcall variant were labeled Dipcall-annotated phaseable, and all other fragments were labeled as having no annotated phaseable site. Because Dipcall reports only high-confidence variants, fragments with no annotated phaseable site may still carry distinguishing variation.

#### Variant Genotyping Performance

To measure how well the genotyping workflow recovers each individual’s variants, we compared vg call genotypes against the truth sets with RTG Tools vcfeval^25^. We used the true positive, false positive, and false negative counts from vcfeval to calculate precision, recall, and F1 score. Results were aggregated across autosomes and reported for all variants, SNPs only, and non-SNP variants only.

Some truth-set variants cannot be recovered by the workflow at any sequencing depth, for example where graph complexity or alignment difficulty prevents reliable genotyping. To separate these from variants missed because of limited coverage, we simulated 120X reads for each GIAB individual and processed them through the WGS pipeline. Truth-set variants that vcfeval called false negatives at this depth were treated as unrecoverable and formed a blacklist for each individual. We excluded blacklisted variants from both the truth set and the call set and repeated the comparison with vcfeval.

#### Variant Phasing Performance

Because SHAPEIT4 phases only input variants that are also present in the reference panel, we evaluated variant phasing at panel-eligible sites only. Using bcftools, we restricted each truth set to heterozygous sites with no missing alleles, then kept only sites matching a panel variant with exact allele matching, using bcftools isec.

We compared SHAPEIT4-phased variants against these truth sets with WhatsHap compare. A switch error occurs when the phase between two adjacent heterozygous sites is reversed relative to the truth. A flip is a single site whose phase is reversed relative to both of its neighbors, which would otherwise count as two switch errors. We report switch error and flip rates for each individual.

To find where heterozygous variants are lost before haplotype inference, we tracked each truth-set heterozygous site through successive stages of the workflow. For each site, we recorded whether it was correctly genotyped, whether it was present in the reference panel, and whether SHAPEIT4 assigned it a phase. The resulting counts show how genotyping accuracy and reference panel representation limit the number of variants available for haplotype inference.

#### Read Phasing Performance

To separate the performance of GRASER phase from errors in haplotype inference, we phased each WGS dataset aligned to the standard HPRCv2 graph against two pairs of paths, the individual’s true haplotype paths from the graph and the personalized paths inferred by GRASER path. To evaluate performance when the target individual is absent from the graph, we also aligned HG002 and HG005 to the HPRCv2-eval graph and phased them against paths inferred by GRASER path.

To compare GRASER with a linear-reference workflow, we aligned each WGS dataset to CHM13 with bwa-mem2^22^. For each individual, we phased the vg call genotypes with WhatsHap phase using these CHM13 alignments, so that both workflows started from the same variants. Reads were then assigned to haplotypes with WhatsHap haplotag.

We defined assignment rate as the proportion of fragments assigned to a haplotype out of all fragments, and calculated it for every dataset and configuration. Because the haplotype of origin is known for simulated reads, reads phased against the true haplotype paths could also be assessed for assignment accuracy, the proportion of assigned fragments placed on the correct haplotype. To evaluate the effect of genomic context, we stratified assignment rate and accuracy by Dipcall annotation for HG002 and HG005, and by difficulty for HG002 only.

Haplotypes inferred by statistical phasing or by WhatsHap are not labeled by parent of origin, and their phase can switch relative to the parental haplotypes along a chromosome or between phase blocks. Assignment accuracy was therefore measured only for reads phased against the true haplotype paths.

### Functional Application: WGBS

#### WGBS Alignment & Read Phasing

The k-mer sampling used to personalize the graph for WGS alignment cannot be applied to WGBS, because bisulfite conversion changes the k-mers in the reads. We therefore built the personalized graph from the individual’s WGS library. Haplotype sampling selects panel haplotypes that match the reads in large blocks, so its paths only approximate the individual’s haplotypes. The paths inferred by GRASER path come from the individual’s own genotypes with statistical phase, and give a closer approximation of the donor haplotypes with longer-range phasing. We therefore replaced the non-reference paths in the diploid personalized graph with the paths inferred by GRASER path. All WGBS libraries were aligned to this graph with methylGrapher using default parameters. For efficiency, each library was split into six chunks with seqkit^23^, aligned separately, and the alignments were concatenated. The aligned reads were then assigned to haplotypes with GRASER phase using default settings, with the paths inferred by GRASER path as targets. Ambiguous reads were not randomly assigned, so that only reads carrying evidence for a haplotype contributed to haplotype-resolved methylation.

#### Allele Specific Methylation

At imprinted loci, DNA methylation differs between the two haplotypes of every individual, because one parental allele is methylated and the other is not. This pattern is set by parent of origin and varies little across tissues and individuals^24^, so imprinted loci provide known allele-specific methylation against which to test GRASER phasing of WGBS reads. We used 68 imprinted DMRs reported by at least two of four studies, as compiled by Akbari et al.^26^, converted from GRCh38 to CHM13 coordinates with the UCSC liftOver utility and the hg38ToHs1 chain file^27^.

Bisulfite conversion leaves methylated cytosines unchanged, so the conversion state recorded in each alignment by methylGrapher gives single-molecule methylation calls^10^. We excluded multimapped reads, then kept alignments with a mapping quality of at least 20 and at least 50 matching bases. Because fastp removes only duplicates with identical sequences, a sequencing error can make a PCR duplicate appear unique, so we also removed duplicates by position. Read pairs whose mates shared 5’ end positions in the graph, soft clipping, and conversion strand were treated as copies of one molecule, and the pair with the highest summed mapping quality was kept. Each inferred haplotype path is an ordered series of graph segments, and concatenating their sequences gives a linear sequence for that haplotype. We converted the methylation calls of each read pair to coordinates on this sequence, a step we call haplotype path linearization. Where both mates covered a CpG, one call was kept if they agreed, and the CpG was dropped from that pair if they disagreed. Read pairs placed on more than one chromosome or spanning more than 10 kb were removed, as were calls on segments the path visits more than once. Methylation calls for each read pair were recorded in modBed format^28^.

Comparing methylation between haplotypes requires the same CpG on both, so we kept CpGs that lie on the same graph segment, in the same orientation, on both haplotype paths, and read CG on both. To limit testing to CpGs with read support for haplotype assignment, we kept CpGs within 150 bp of a heterozygous variant. The heterozygous variants within this window had to be either all phased or a single unphased variant. Phased variants whose 150 bp windows overlapped formed one phase unit, and each unphased variant formed its own. Read pairs with calls on both strands of a haplotype path were removed. A C/T or G/A SNV cannot be distinguished from bisulfite conversion on one strand. A read pair crossing such a variant was therefore kept only if it also crossed a variant visible on its strand, or an indel, MNV, or SV. Each CpG contributed one call per read pair. chrX was excluded because the donor is male.

For each tissue, we compared methylation between haplotypes with DSS, which uses neighboring CpGs to estimate variation in the absence of replicates^29^. CpGs with calls on both haplotypes were tested with DSS 2.58.0 DMLtest, using a smoothing span of 200 bp. Phase units were laid end to end, 10 kb apart, so that smoothing never combined CpGs from different units. The false discovery rate was controlled at 5% per tissue across all tested CpGs with the Benjamini-Hochberg procedure. Significant CpGs were grouped into regions with the DSS callDMR procedure, merging regions within 50 bp. Each region had to be longer than 50 bp, with more than 3 tested CpGs, more than half of them significant. Regions were split at gaps between tested CpGs larger than 100 bp, and each piece was rechecked against the same criteria. The methylation difference of a region was the mean of DSS’s smoothed haplotype 1 minus haplotype 2 methylation across its tested CpGs. We kept regions with an absolute difference of at least 0.2 and at least 3 distinct read pairs on each haplotype. Regions whose depth exceeded 10 times the median depth of the tissue’s regions passing the difference threshold were removed.

An imprinted region was considered detected in a tissue when a tested CpG of a DMR fell within it. Haplotype labels from statistical phasing do not indicate parent of origin. Within one individual, however, an imprinted region should have the same haplotype methylated in every tissue, so we checked for that consistency.

## RESULTS

Because the haplotype paths inferred by GRASER path depend on graph-based genotyping and statistical phasing, we first evaluated both steps as applied in our workflow.

### Genotyping Performance

We compared variants called by vg call with the graph-derived truth sets using RTG Tools vcfeval (Fig. 2). Genotyping performance was consistent across individuals, with F1 scores of approximately 92 to 94% for the GIAB cohort on the standard HPRCv2 graph, and a mean F1 score of 92.3% for the 1kGP cohort (Fig. 2C, D). Recall ranged from 93.99 to 95.81%, consistently higher than precision, ranging from 89.00 to 91.64%, so most truth-set variants were recovered, but a subset of called variants did not match the truth set (Fig. 2A, B). Simulated reads performed slightly better than real sequencing data, but the difference was small.

**Figure 1.**
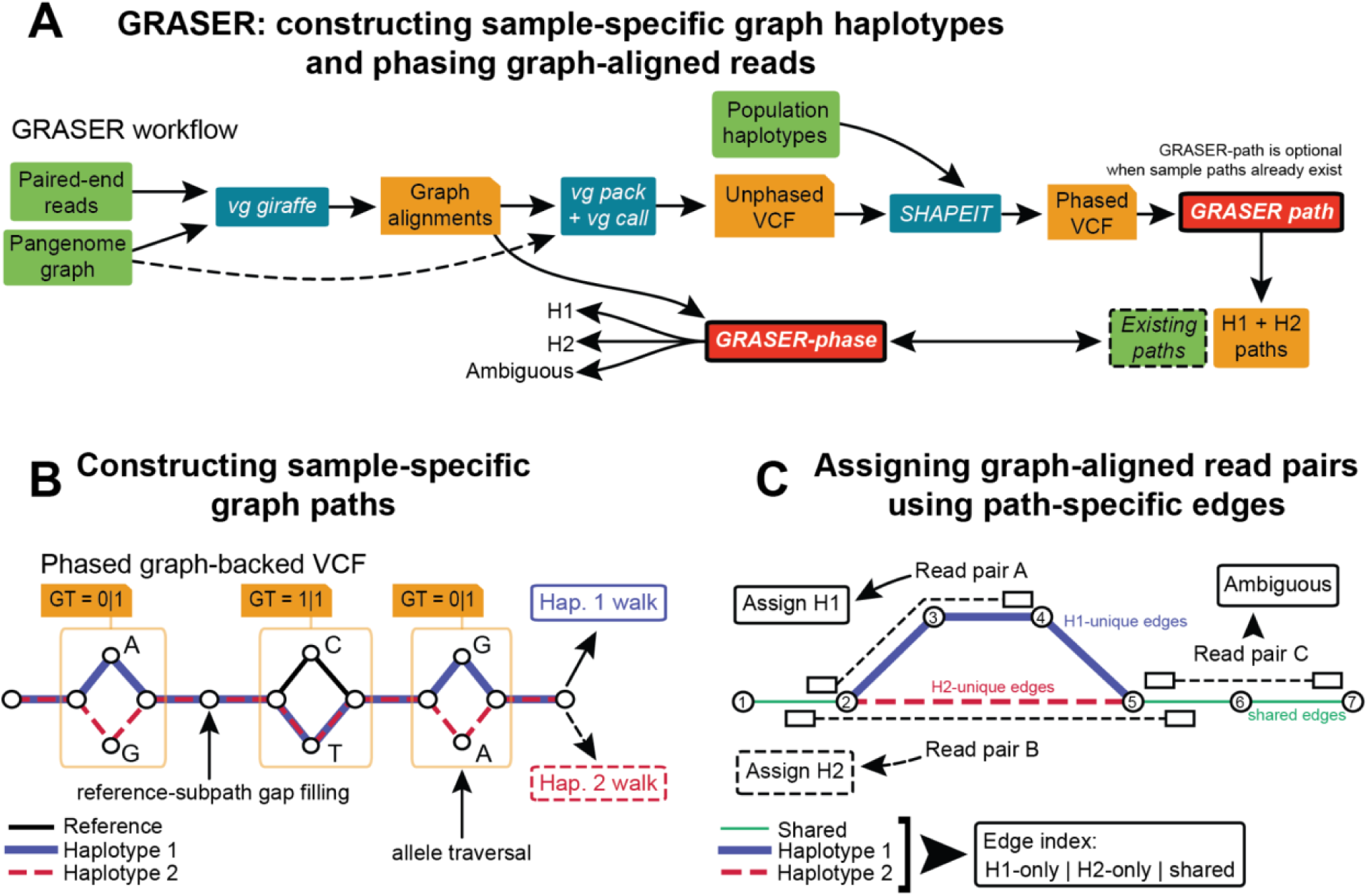
GRASER infers sample-specific haplotype paths through a pangenome graph and assigns graph-aligned reads to them. **A.** GRASER workflow. Green boxes are inputs, teal boxes are external tools, orange boxes are intermediate files, and red boxes are GRASER modules. Paired-end reads are aligned to the pangenome graph with vg giraffe, and variants are genotyped from these alignments with vg pack and vg call against the complete graph (dashed arrow). The resulting VCF is phased with SHAPEIT4 using population haplotypes. GRASER path builds two haplotype paths (H1 and H2) from the phased VCF. This step can be skipped when sample paths already exist in the graph (dashed green box). GRASER phase then assigns each aligned read pair to H1 or H2, or labels it ambiguous. **B.** Construction of sample-specific graph paths. Nodes (circles) contain sequence, and edges connect adjacent sequences. Black, blue, and dashed red lines mark the reference, haplotype 1, and haplotype 2 paths. Three variable sites (orange boxes) are shown with their phased genotypes from the graph-backed VCF and the base carried by each allele node. Each allele corresponds to a traversal through its site, and at the homozygous site (1|1) both haplotypes take the non-reference allele. Allele traversals are chained together, and gaps between sites are filled with the reference subpath, giving one contiguous walk per haplotype. **C.** Assignment of read pairs using path-specific edges. The edges of the two haplotype paths are indexed as H1-only (blue), H2-only (dashed red), or shared (green), and haplotype-unique edges in a read pair’s alignment are evidence for that haplotype. Here, the H1 path is 1>2>3>4>5>6>7 and the H2 path is 1>2>5>6>7. Read pairs are drawn as two boxes joined by a dashed line. Read pair A traverses edges 2>3 and 3>4, which are unique to H1, and is assigned to H1. Read pair B traverses edge 2>5, which is unique to H2, and is assigned to H2. Read pair C traverses only shared edges, so it carries no haplotype-distinguishing information and is labeled ambiguous.

**Figure 2.**
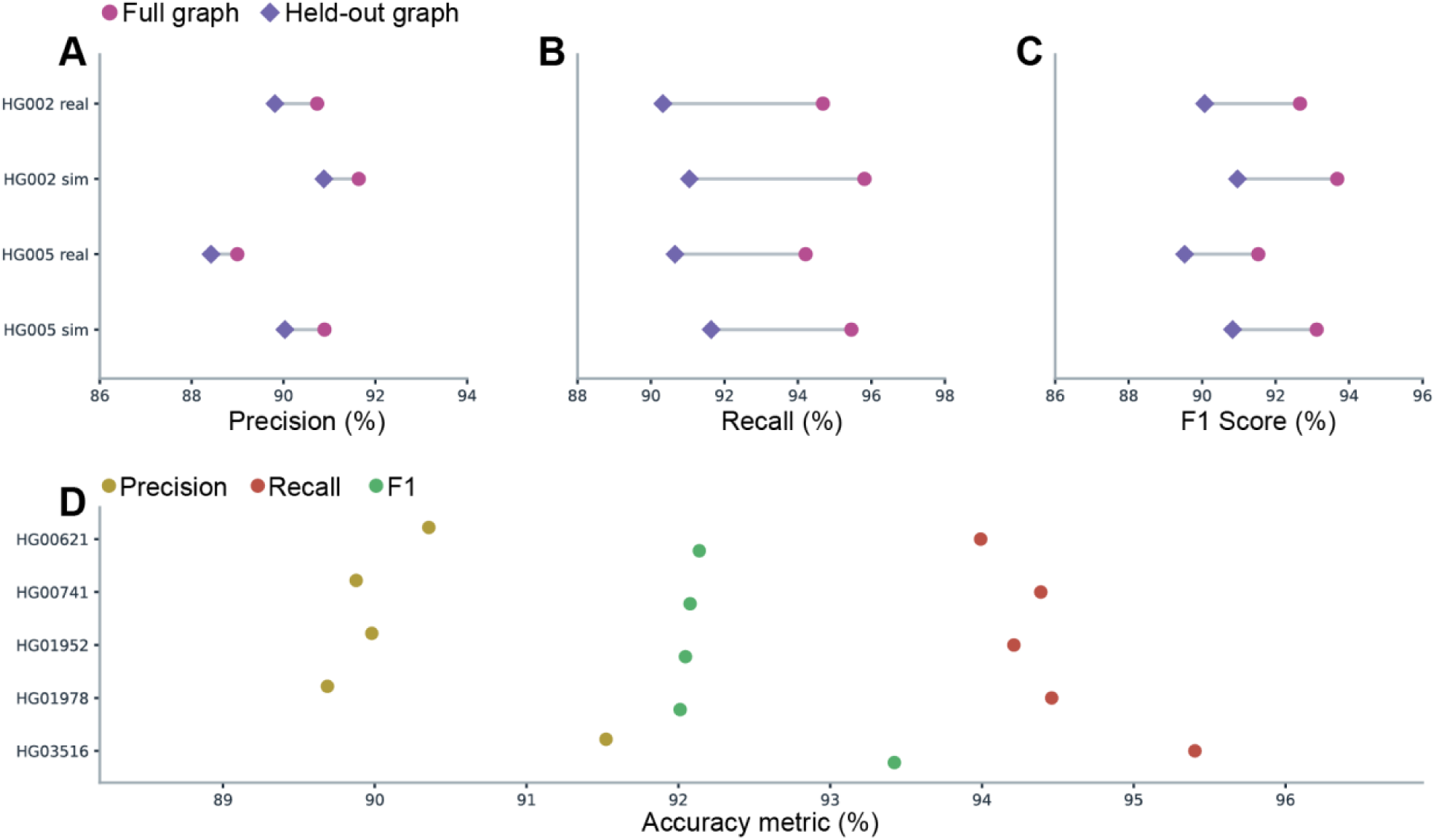
Graph-based genotyping recovers most truth-set variants, with lower precision than recall and a modest loss when the individual is excluded from the graph. Genotypes from vg call were compared with each individual’s graph-derived truth set using RTG Tools vcfeval, and precision, recall, and F1 score were calculated from true positive, false positive, and false negative counts pooled across autosomes. **A–C**. Precision (A), recall (B), and F1 score (C) for HG002 and HG005, from real and simulated WGS aligned to the standard HPRCv2 graph (magenta circles) or the HPRCv2-eval graph (purple diamonds), which excludes both individuals. Grey lines connect the two graphs for each dataset. **D**. Precision (gold), recall (red), and F1 score (green) for the five 1kGP individuals, from real WGS aligned to the standard HPRCv2 graph.

Excluding the evaluated individual’s assemblies from the reference graph modestly reduced genotyping performance (Fig. 2). For HG002 and HG005, F1 scores decreased by an average of ∼2.4 percentage points, mainly through reduced recall rather than precision. This reduction was larger for non-SNP variants than for SNPs. Variants that remained false negatives even at 120X simulated coverage made up approximately 4 to 4.5% of the GIAB truth sets. Excluding these variants increased F1 scores by an average of ∼2.4 percentage points, so a portion of missed variants cannot be recovered by the workflow at any depth.

### Variant Phasing Performance

We next evaluated whether the workflow produces enough correctly phased variants to construct haplotype paths (Fig. 3). Only approximately 74 to 76% of truth-set heterozygous variants were present in the SHAPEIT4 reference panel, so about a quarter could not be phased by this approach. Overall, 70.9 to 73.9% of truth-set heterozygous sites were correctly genotyped, present in the panel, and phased (Fig. 3A). Of the panel-eligible variants, approximately 99% were correctly genotyped using the standard HPRCv2 graph, with a modest reduction using the HPRCv2-eval graph (Fig. 3B). SHAPEIT4 assigned a phase to nearly all correctly genotyped, panel-eligible heterozygous variants.

**Figure 3.**
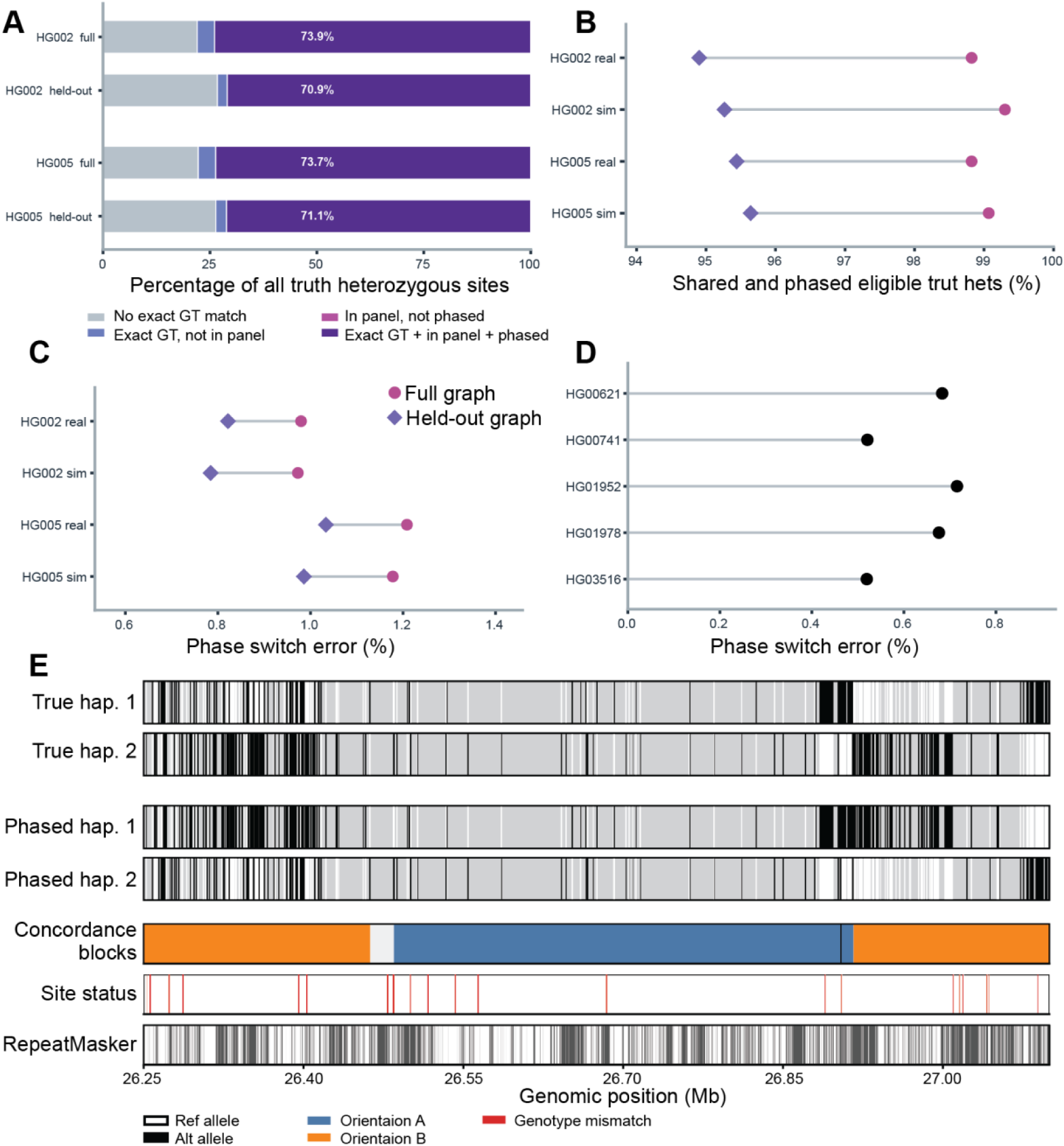
Most heterozygous variants that are correctly genotyped and present in the reference panel are phased, with low switch error. **A.** Fate of truth-set heterozygous variants through genotyping and statistical phasing, for HG002 and HG005 simulated WGS aligned to the HPRCv2 graph (“full”) or the HPRCv2-eval graph (“held-out”). Bars partition all truth heterozygous sites into four mutually exclusive categories: no exact genotype match (grey), correct genotype but absent from the reference panel (light blue), correct genotype and in the panel but not phased (pink), and correct genotype, in the panel, and phased (purple, percentage shown). **B**. Recovery of panel-eligible heterozygotes, the proportion of heterozygous sites in the panel-restricted truth set that were shared with and phased in the SHAPEIT4 output, for real and simulated WGS on the HPRCv2 (magenta circles) and HPRCv2-eval (purple diamonds) graphs. **C.** Switch error rates for HG002 and HG005, computed with WhatsHap compare against the graph-derived truth sets, with the same symbols as B. **D.** Switch error rates for the five 1kGP individuals, from real WGS on the HPRCv2 graph. These individuals are in the reference panel. **E.** Local phase concordance for HG002 simulated WGS aligned to the HPRCv2 graph, on CHM13 chromosome 1 (26.25 to 27.10 Mb). Truth-set heterozygous sites were matched to the SHAPEIT4 output by variant ID. Sites with matching heterozygous genotypes were classified by whether the SHAPEIT4 haplotype order agreed with the truth (orientation A, blue) or was reversed (orientation B, orange). The top four tracks show true and SHAPEIT4-phased haplotypes at these sites, with reference (white) and alternate (black) alleles. Consecutive sites with the same orientation were grouped into concordance blocks, so each change between blocks marks a switch error. Sites more than 1 Mb apart were segmented separately. Short discordant runs of up to 3 sites within 10 kb, flanked by at least 5 concordant sites on each side, were treated as isolated errors (black markers) rather than block boundaries. Concordance blocks describe local agreement with the truth haplotypes and are distinct from the phase blocks reported by SHAPEIT4. Site status marks genotype mismatches (red). The bottom track shows RepeatMasker annotations.

Switch error rates were low for all seven individuals, ranging from approximately 0.5% to 1.2% (Fig. 3C, D). The GIAB individuals had slightly lower switch error rates on the HPRCv2-eval graph, despite fewer eligible variants being recovered (Fig. 3C). The 1kGP individuals generally had lower switch error rates than the GIAB individuals, consistent with their inclusion in the reference panel (Fig. 3D). An example interval shows the relationship between truth-set heterozygous variants, variants recovered by genotyping, and the haplotype blocks inferred by statistical phasing (Fig. 3E).

### Read Phasing Performance

To test GRASER phase directly, we first phased simulated reads against the true parental haplotype paths from the standard HPRCv2 graph. Under these conditions, GRASER assigned 24.5% and 22.6% of fragments to a haplotype for HG002 and HG005, respectively (Fig. 4A). Fragments overlapping high-confidence heterozygous variants were assigned at substantially higher rates than other fragments, while maintaining high accuracy. For HG002, these fragments were assigned at a rate of 93.1%, with an accuracy of 98.4%; for HG005, assignment was 94.3% and accuracy was 98.2%. Conversely, fragments that did not overlap such variants had assignment and accuracy rates of 6.9% and 91.5% in HG002, and 6.1% and 89.6% in HG005.

**Figure 4.**
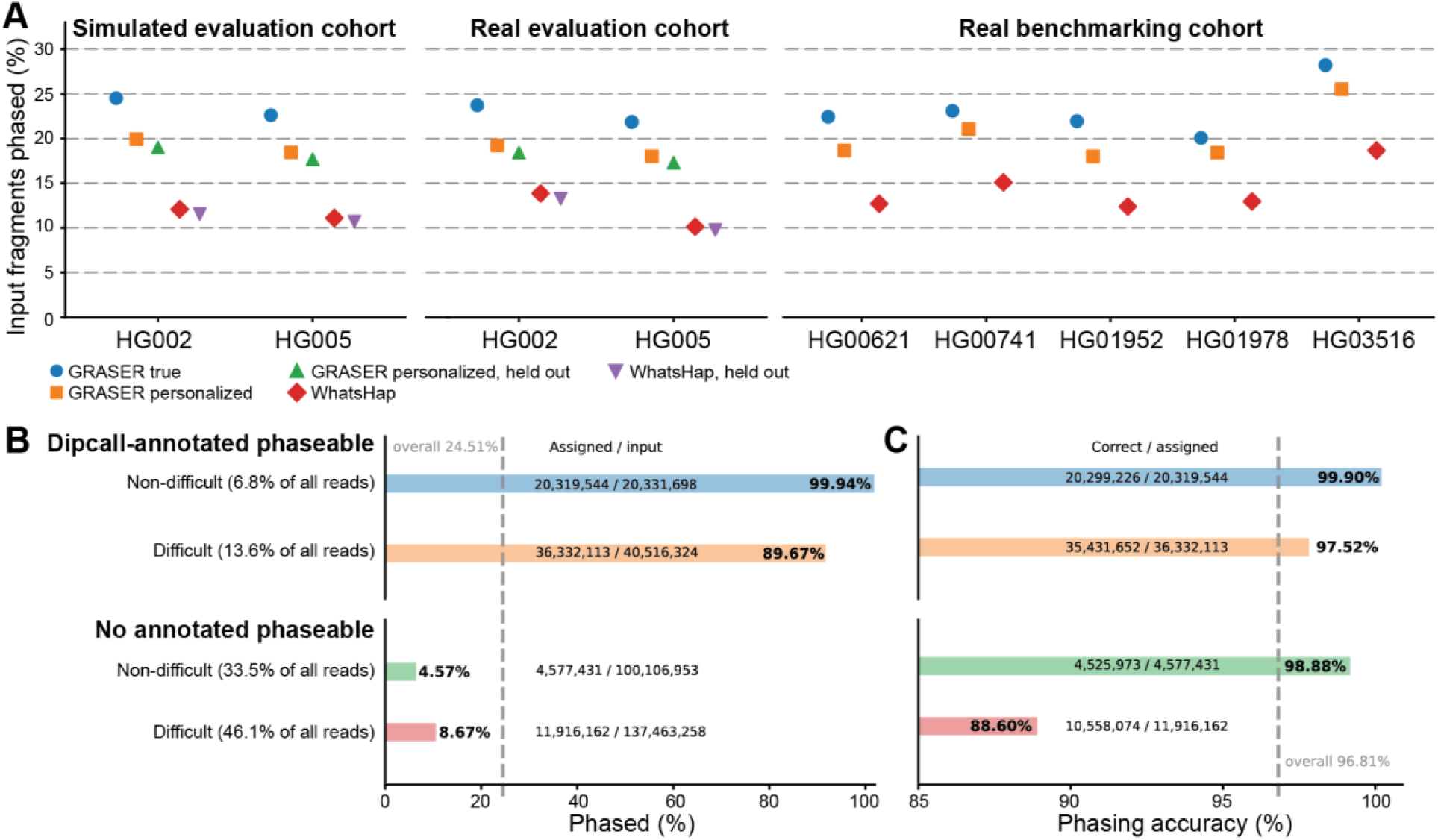
GRASER assigns more fragments than a linear-reference workflow, and assigns fragments carrying distinguishing variation with high accuracy. **A**. Fragment assignment rate, the percentage of input fragments assigned to a haplotype, for every configuration. Results are shown for HG002 and HG005 simulated WGS, HG002 and HG005 real WGS, and real WGS from the five 1kGP individuals. GRASER phase was run against the individual’s true haplotype paths in the HPRCv2 graph (blue circles), and against haplotype paths inferred by GRASER path from alignments to the HPRCv2 graph (orange squares) or the HPRCv2-eval graph (green triangles). WhatsHap phasing of CHM13 alignments is shown using vg call genotypes from alignments to the HPRCv2 graph (red diamonds) or the HPRCv2-eval graph (purple triangles). **B, C.** HG002 simulated WGS phased against the true haplotype paths in the HPRCv2 graph. Fragments are stratified by whether at least one mate overlaps a Dipcall heterozygous variant (Dipcall-annotated phaseable) or not (no annotated phaseable site), and by overlap with GIAB difficult regions. Because Dipcall reports only high-confidence variants, fragments without an annotated phaseable site can still be phased. Each category’s share of input fragments is given beside its label. B shows the percentage of input fragments assigned to a haplotype, and C shows the percentage of assigned fragments placed on the correct haplotype, with numerators and denominators on each bar. Dashed lines mark the overall assignment rate (24.51%) and accuracy (96.81%). The x-axis in C starts at 85%.

To examine how genomic context affects read phasing, we stratified HG002 fragments by Dipcall annotation and by overlap with GIAB difficult regions (Fig. 4B, C). A fragment was considered Dipcall-annotated phaseable if at least one mate overlapped a high-confidence heterozygous variant between the parental assemblies (Methods). Overall, more than 96% of assigned fragments were placed on the correct parental haplotype for both HG002 and HG005 (Fig. 4C). Dipcall-annotated phaseable fragments outside difficult regions performed best, with 99.9% assigned and 99.9% of assigned fragments correct. Dipcall-annotated phaseable fragments in difficult regions were assigned at a lower rate (89.7%) but remained highly accurate (97.5%). Fragments with no phaseable site annotation and outside difficult regions were assigned at a rate of 4.6%, with an accuracy of 98.9%. Fragments with no phaseable site annotation and within difficult regions were assigned at a rate of 8.7% with an accuracy of 88.6%. Overall, fragments with no annotated phaseable site were assigned much less often, and those in difficult regions had the lowest accuracy. GRASER therefore assigns fragments carrying distinguishing variation with high accuracy, and its performance decreases in difficult regions.

We next phased reads against the haplotype paths inferred by GRASER path (Fig. 4A). On the standard HPRCv2 graph, inferred paths reduced the fragment assignment rate by approximately 4 percentage points relative to the true parental paths. Inferred paths lack variants missed by genotyping or absent from the reference panel, and can carry phasing errors, so fewer reads traverse a haplotype-unique edge. With inferred paths, aligning to the HPRCv2-eval graph instead of the standard graph reduced the assignment rate by less than 1 additional percentage point on average, so performance depends little on whether the individual is in the graph. The loss from haplotype inference was therefore much larger than the loss from excluding the individual from the graph. Genotyping recovered about 99% of panel-eligible variants, but a quarter of heterozygous variants were absent from the reference panel, pointing to the panel as the main limitation (Fig. 3B). These trends were consistent between simulated and real sequencing data.

To compare GRASER with a linear-reference workflow, we phased the same WGS datasets with WhatsHap, starting from the same vg call variants (Fig. 4A). GRASER assigned a greater proportion of fragments than WhatsHap in every matched configuration. For the five 1kGP individuals, GRASER assigned an average of 20.3% of fragments using inferred haplotype paths, compared with 14.4% for WhatsHap. GRASER also assigned more fragments than WhatsHap for the GIAB individuals, including on the HPRCv2-eval graph.

GRASER phase completed in approximately two hours per sample (mean 1.83 ± 0.17), using six threads and less than 40 GB of memory.

## WGBS

To test whether GRASER-phased WGBS reads recover allele-specific methylation, we examined imprinted loci, where one parental allele is methylated and the other is not. Imprinted methylation is set by parent of origin and generally varies little across tissues and individuals^24^, so we expected most of these loci to show allele-specific methylation across the donor’s tissues. Of the 68 imprinted regions, 38 contained at least one tested CpG in at least one tissue, and the other 30 had no CpG shared by both haplotypes near a heterozygous variant. Of the 38 regions, 24 had a DMR in at least one tissue, and 8 had a DMR in all six tissues. All 22 regions with DMRs in two or more tissues had the same haplotype methylated in every tissue, consistent with the usual stability of imprinted methylation across tissues.

Between the 3’ end of GPR1-AS and the ZDBF2 promoter, a known imprinted DMR showed allele-specific methylation in all six tissues (Fig. 5). Haplotype 1 was hypomethylated relative to haplotype 2 in every tissue. In thyroid, the DMR contained 13 tested CpGs, covered by 31 and 41 read pairs on haplotypes 1 and 2, with haplotype 2 more methylated by 0.79.

**Figure 5.**
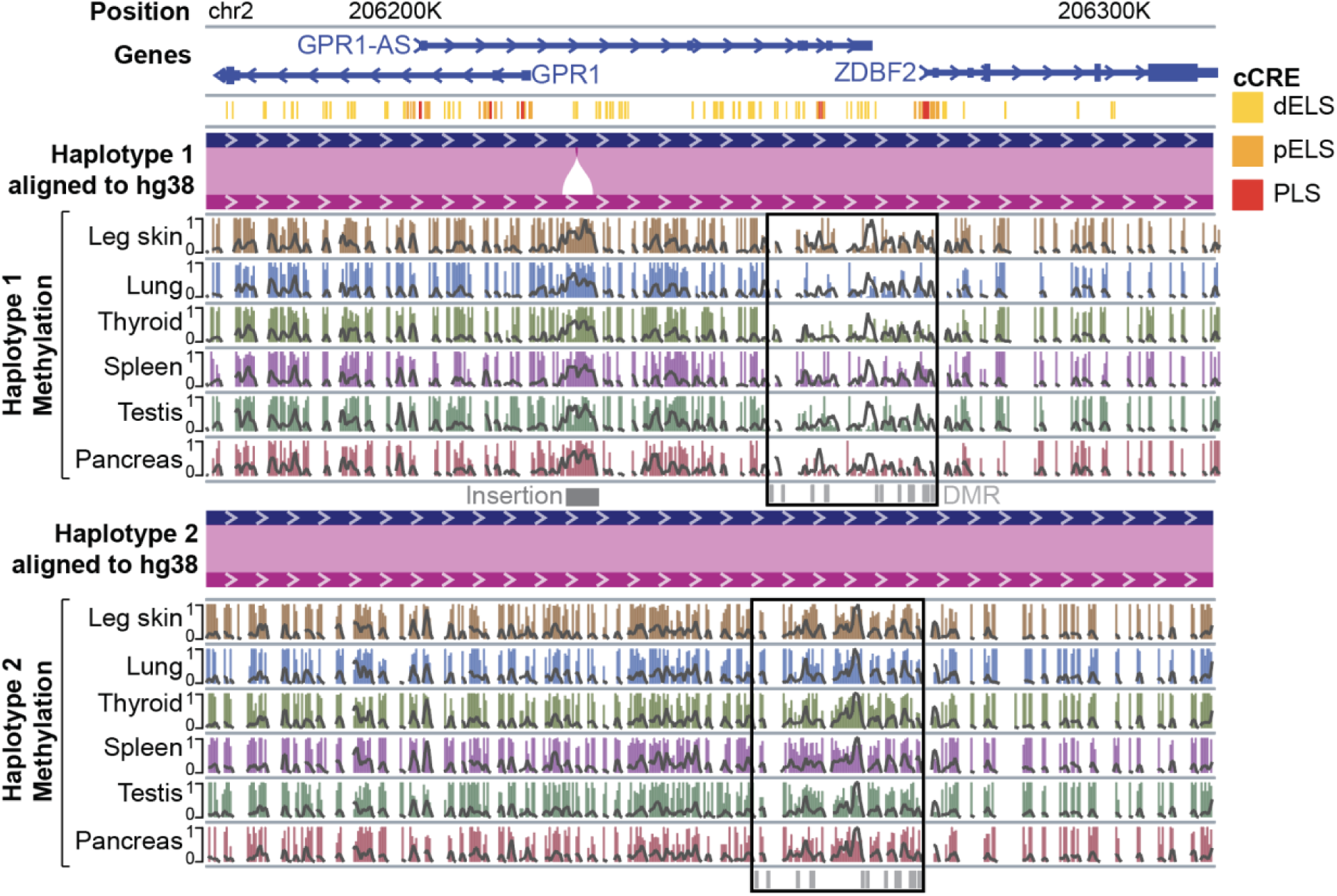
GRASER-phased WGBS reads recover allele-specific methylation at the ZDBF2 imprinted DMR and measure methylation within a haplotype-specific insertion. WGBS from six tissues of one EN-TEx donor were aligned with methylGrapher to a graph carrying the haplotype paths inferred by GRASER path from the donor’s WGS, then assigned to haplotypes with GRASER phase. The view spans GPR1, GPR1-AS, and ZDBF2 on chromosome 2, shown in the WashU Epigenome Browser^30^ with both haplotypes projected onto hg38. Tracks show gene annotations, then ENCODE candidate cis-regulatory elements (cCREs: distal enhancer-like, dELS, yellow; proximal enhancer-like, pELS, orange; promoter-like, PLS, red), then each haplotype’s alignment to hg38, then methylation in each tissue on haplotype 1 (upper six tracks) and haplotype 2 (lower six tracks). Bars show CpG methylation (0 to 1) and grey lines show relative coverage, both from reads assigned to that haplotype. Black boxes mark the imprinted DMR between the 3’ end of GPR1-AS and the ZDBF2 promoter, where haplotype 1 is hypomethylated relative to haplotype 2 in every tissue. The grey bar marks an insertion on haplotype 1 within a candidate enhancer, whose sequence is highly methylated on haplotype 1 in every tissue.

Graph alignment also allows methylation to be measured in sequence absent from the reference. Upstream of the ZDBF2 imprinted DMR, haplotype 1 carries an insertion within a candidate enhancer (Fig. 5). Reads from the inserted sequence have no position on the reference path, so surjection would discard them. Because these reads were aligned and phased on the graph, we could see that the insertion places a block of highly methylated sequence within the enhancer on haplotype 1, in every tissue. GRASER therefore enables haplotype-resolved profiling of DNA methylation within structural variants from short-read WGBS data.

## DISCUSSION

Here, we present GRASER, which phases graph-aligned short reads against haplotype paths inferred for each individual, using established graph genotyping and statistical phasing to build those paths. Because these paths come from the individual’s own reads, GRASER does not require the individual’s assemblies, either in the graph or in advance. GRASER phase uses edges unique to one haplotype path as evidence, so reads are phased directly from graph alignments without conversion to a linear reference. Against true parental haplotype paths, GRASER assigned 22.6 to 24.5% of simulated fragments to a haplotype, and more than 96% of those were placed on the correct haplotype. Inferred haplotype paths reduced the assignment rate by about 4 percentage points. GRASER assigned more fragments than a WhatsHap-based linear-reference workflow in every configuration tested. Applied to WGBS, GRASER recovered allele-specific methylation at imprinted loci and measured haplotype-resolved methylation within a structural variant.

In simulated WGS, read phasing depended on whether a fragment carried distinguishing variation and on the genomic context it came from. Dipcall-annotated phaseable fragments outside GIAB difficult regions were nearly all assigned, and nearly all were assigned correctly, at 99.9% for both. This follows from edge-based assignment, because variation between haplotypes creates edges unique to one of them, and a fragment that crosses such an edge carries direct evidence for its haplotype.

In simulated WGS, phasing was less effective for fragments in difficult regions and for fragments with no annotated phaseable site. Dipcall reports variants only where the two parental assemblies can be aligned with confident synteny, so heterozygous variants in duplicated regions are excluded. As a result, some fragments with no annotated phaseable site still cross a haplotype-unique edge and are assigned correctly. In difficult regions, incorrect alignments can traverse edges unique to the wrong haplotype, which lowers accuracy. Phasing performance therefore depends on both the availability of distinguishing variation and the reliability of graph alignment.

Inferred haplotype paths depend on graph genotyping and statistical phasing, and errors at either step carry into the paths. Graph genotyping recovered most truth-set variants but had lower precision than recall, which may reflect alignment errors, the genotyping algorithm, or differences in how variants are represented. About 4 to 4.5% of truth-set variants were missed even at 120X coverage, so their loss cannot come from a lack of data. At these sites, the workflow either cannot place reads correctly in the graph or cannot genotype from them, so they are inaccessible to this approach and are appropriately excluded from evaluation. Because the blacklist comes from the same pipeline, performance after excluding it describes variants this workflow can recover, not an independent estimate.

The larger limitation lies in statistical phasing. About a quarter of truth-set heterozygous variants were absent from the SHAPEIT4 reference panel and could not be phased, while nearly all panel-eligible variants were genotyped and phased with low switch error. The panel was built from short-read calls, so it likely under-represents indels and structural variants, which are also the variants most often represented differently in the graph and in the panel. GRASER path assigns a random phase to variants that SHAPEIT4 leaves unphased, which can introduce switch errors into the inferred paths. Larger short-read panels would add common variants but still miss many structural variants that pangenome genotyping recovers. A more promising route is to genotype a large cohort against the pangenome and phase it as a cohort, so that phasing draws on the same variant representation as the graph and needs no external panel.

Pangenome methods are often evaluated on individuals whose assemblies are in the graph, which can overstate performance on new individuals. Excluding HG002 and HG005 from the graph reduced genotyping recall, but reduced the assignment rate by less than 1 percentage point. This works because most of a new individual’s variation is common, and the 460 haplotypes of the second HPRC release capture over 99% of common variation in the All of Us cohort^9^. As pangenomes become more complete for common variation, the haplotype paths GRASER infers for new individuals should approach those of individuals in the graph. The comparison shows that assignment rate is stable across graphs, but does not establish that accuracy is.

GRASER assigned a greater proportion of sequencing fragments than the conventional WhatsHap-based workflow across all evaluated conditions. The higher assignment rate most likely comes from how variants are phased. Read-based phasing with WhatsHap links two heterozygous variants only when a read or read pair spans both, so variants farther apart than a fragment cannot be phased, and reads at isolated variants cannot be assigned. Statistical phasing imposes no such distance constraint, so nearly all panel-eligible variants are placed on chromosome-length haplotype paths. Graph alignment may add to this by retaining reads carrying non-reference alleles that linear alignment can lose to reference bias. Because these experiments compare complete workflows, they do not separate these contributions. Since accuracy was not compared, the results show an advantage in assignment rate, not in accuracy.

GRASER recovered allele-specific methylation at known imprinted loci from short-read WGBS, with the same haplotype methylated across tissues. Because reads are aligned and phased on the graph, methylation can also be measured within structural variants on the haplotype that carries them, which linear-reference workflows cannot do. Our analysis tested only CpGs present on both haplotypes, so CpGs created or destroyed by a heterozygous variant, among the most direct cis effects on methylation, were not tested.

Accuracy was measured on error-free simulated reads phased against true parental paths, conditions more favorable than real data. The genotyping and phasing truth sets come from the same pangenome used for alignment, so they are not fully independent of it. The 1kGP individuals are in the SHAPEIT4 reference panel, which likely contributed to their lower switch error rates.

By default, GRASER weights every haplotype-unique edge equally. Weights could instead reflect the sequence an edge spans, down-weighting edges whose composition or low complexity makes alignment unreliable. They could also be tuned to the data type, because bisulfite conversion reduces sequence complexity, so an edge reliable for WGS reads may be ambiguous for converted reads. Longer reads would cross more distinguishing edges per fragment, increasing the share of reads GRASER can assign.

GRASER connects the haplotype information in pangenome graphs to individual sequencing reads, without requiring an individual’s assemblies. Because reads are assigned on the graph, haplotype-resolved measurements extend to sequence absent from the linear reference, including structural variants. Once reads are assigned to haplotype paths, each path can be linearized, so the data can be represented on two linear haplotype sequences. Existing tools built for linear references can then analyze these data directly, without the loss of non-reference reads that surjection to a fixed reference causes. Choosing a genome for analysis still involves important considerations^6^, but this removes a logistical barrier to using pangenome references within existing workflows, particularly for non-specialists. With that barrier lowered, the human pangenome can move from a specialist resource into routine analysis, carrying its representation of human genetic diversity into everyday studies of molecular phenotypes.

## DATA & CODE AVAILABILITY

GRASER is available at https://github.com/twlab/GRASER. The deterministic lattice read simulator is available in MoRGANA (https://github.com/juanfmacias/MoRGANA). HPRC release 2 assemblies are available at https://humanpangenome.org/data/, and the release 2 Minigraph-Cactus graphs at https://github.com/human-pangenomics/hpp_pangenome_resources.

